# The assembled and annotated genome of the pigeon louse *Columbicola columbae*, a model ectoparasite

**DOI:** 10.1101/2020.10.08.330514

**Authors:** James G. Baldwin-Brown, Scott M. Villa, Anna I. Vickrey, Kevin P. Johnson, Sarah E. Bush, Dale H. Clayton, Michael D. Shapiro

**Affiliations:** School of Biological Sciences, University of Utah, Salt Lake City, UT, USA; Department of Biology, O. Wayne Rollins Research Center, Emory University, Atlanta, GA; Illinois Natural History Survey, Prairie Research Institute, University of Illinois, Champaign, Illinois, USA 61820

**Keywords:** Genome assembly, Genome annotation, Insect genomics, Ectoparasitism, Phthiraptera, Ischnocera

## Abstract

The pigeon louse *Columbicola columbae* is a longstanding and important model for studies of ectoparasitism and host-parasite coevolution. However, a deeper understanding of its evolution and capacity for rapid adaptation is limited by a lack of genomic resources. Here, we present a high-quality draft assembly of the *C. columbae* genome, produced using a combination of Oxford Nanopore, Illumina, and Hi-C technologies. The final assembly is 208 Mb in length, with 12 chromosome-size scaffolds representing 98.1% of the assembly. For gene model prediction, we used a novel clustering method (*wavy_choose*) for Oxford Nanopore RNA-seq reads to feed into the *MAKER* annotation pipeline. High recovery of conserved single-copy orthologs (BUSCOs) suggests that our assembly and annotation are both highly complete and highly accurate. Consistent with the results of the only other assembled louse genome, *Pediculus humanus*, we find that *C. columbae* has a relatively low density of repetitive elements, the majority of which are DNA transposons. Also similar to *P. humanus*, we find a reduced number of genes encoding opsins, G protein-coupled receptors, odorant receptors, insulin signaling pathway components, and detoxification proteins in the *C. columbae* genome, relative to other insects. We propose that such losses might characterize the genomes of obligate, permanent ectoparasites with predictable habitats, limited foraging complexity, and simple dietary regimes. The sequencing and analysis for this genome were relatively low-cost, and took advantage of a new clustering technique for Oxford Nanopore RNAseq reads that will be useful to future genome projects.

## Introduction

Parasites represent a large proportion of eukaryotic biodiversity, and it is estimated that 40% of insect diversity is parasitic (de Meeûs and Renaud 2002). Parasitic lice (Insecta: Phthiraptera) comprise a group of about 5000 species that parasitize all orders of birds and most orders of mammals (Mullen and Durden 2009; Clayton *et al*. 2015). Two thirds of louse species are associated with only a single host species (Durden and Musser 1994; Smith 2004). The genus *Columbicola* comprises 91 known species, all found on pigeons or doves (Bush *et al*. 2009; Gustafsson *et al*. 2015; Adly *et al*. 2019); most of these louse species are found on a single host species (Johnson *et al*. 2007, 2009).

Like all feather lice (suborder Ischnocera), *Columbicola* are “permanent” parasites that complete their entire life cycle on the body of the host (Marshall 1981). Feather lice feed primarily on feathers, which they metabolize with the assistance of endosymbiotic bacteria (Fukatsu *et al*. 2007). The feather damage caused by lice has a chronic effect that leads to reduced host survival (Clayton *et al*. 1999) and mating success (Clayton 1990). Birds are able to defend themselves against feather lice by preening them with the beak. However, *Columbicola* lice escape from preening by hiding in grooves between feather barbs, and the sizes of these grooves scale with host body size. In micro-evolutionary time, the result is stabilizing selection on body size of lice (Clayton *et al*. 1999; Bush and Clayton 2006). In macro-evolutionary time, the result is that host defense (preening) and body size interact to reinforce the host specificity and size matching of *Columbicola* species to their hosts (Clayton *et al*. 2003). Similarly, selection for visual crypsis drives the evolution of color similarities between *Columbicola* species their hosts (Bush *et al*. 2010, 2019).

Within the feather lice, the biology of *C. columbae* (Fig. 1) is better known than that of any other louse species, including details about its morphology, physiology, ecology, and behavior (Martin 1934; Stenram 1956; Rakshpal 1959; Nelson and Murray 1971; Rudolph 1983; Clayton 1990, 1991; Clayton and Tompkins 1995; Clayton *et al*. 1999, 2003; Bush *et al*. 2006; Bush and Clayton 2006; Clayton *et al*. 2008; Harbison and Clayton 2011). A unique feature of the *C. columbae* study system is that its host, the rock pigeon *Columba livia*, has been under artificial selection by pigeon breeders for millennia, resulting in dramatic phenotypic variation (Darwin 1868; Shapiro and Domyan 2013), similar to that seen across the 300+ other species of pigeons and doves (Gibbs *et al*. 2001). This variation makes it possible to transfer *C. columbae* among diverse size and color phenotypes within the single native host species. Recently, we showed that switching lice to pigeons of different sizes and colors elicits rapid population-level changes in louse size and color (Bush *et al*. 2019; Villa *et al*. 2019). Despite the wealth of phenotypic data about real-time adaptation of *C. columbae* to changes in host environment, the underlying molecular mechanisms remain unknown.

**Fig. 1.**
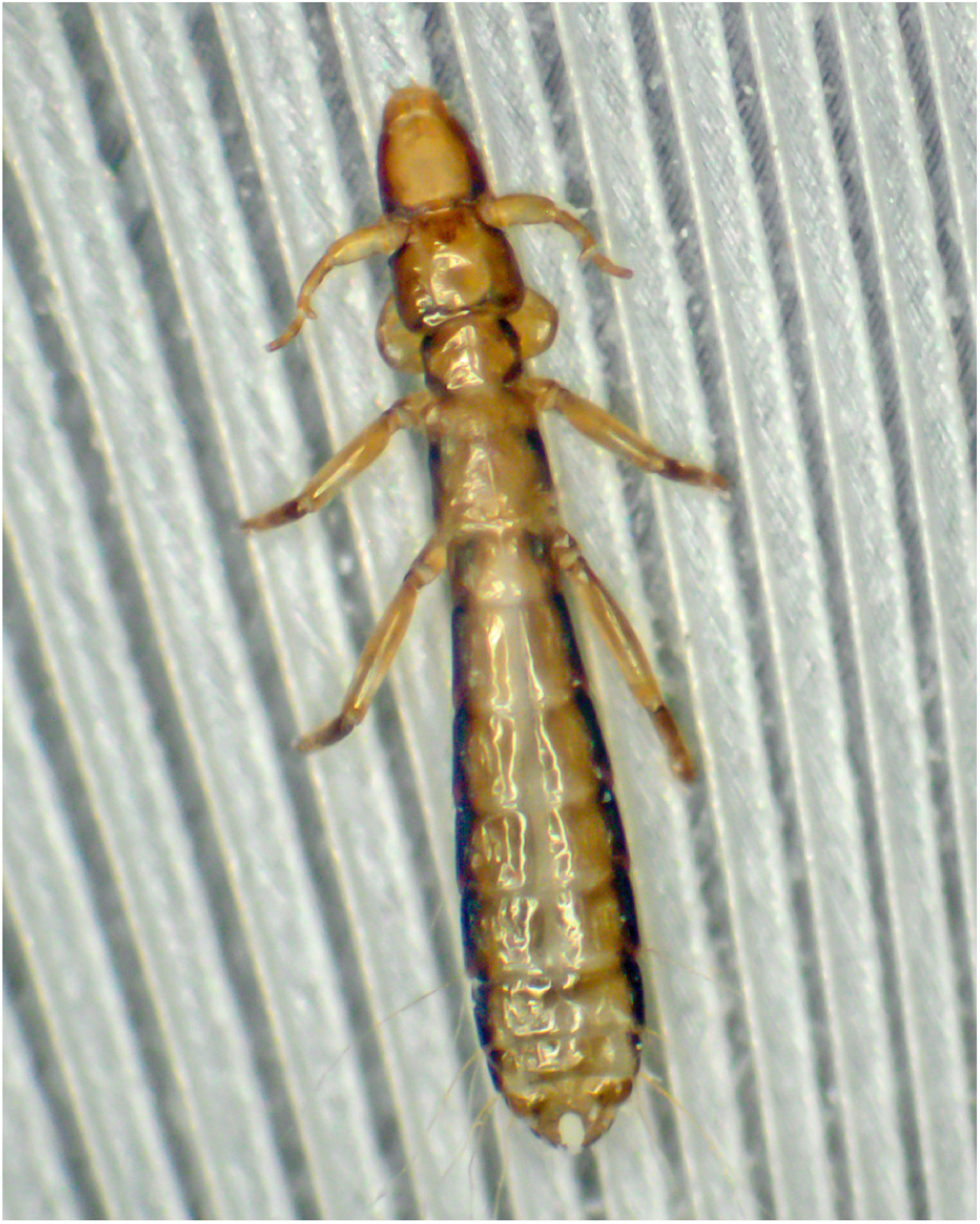
Male *Columbicola columbae* clinging to a rock pigeon feather. The thumblike processes on the antennae are used to grasp a female when mating. Photo by Scott Villa and Juan Altuna.

A deeper understanding of louse evolution and genetics is limited largely by a paucity of genomic resources. The louse with the best available genomic resources is the blood-feeding human body louse *Pediculus humanus*, the draft genome of which was assembled using low-coverage shotgun sequencing (Kirkness *et al*. 2010). *P. humanus* had the smallest insect genome known at that time (108 Mb), with a repertoire of 10,773 annotated genes. Presently, what we know about the genomic signatures of parasitism in Phthiraptera is largely limited to this one species.

Here, we report a high-quality draft genome assembly and annotation for *C. columbae* that incorporates short-read Illumina (Bennett 2004) sequences, long-read Oxford Nanopore (Jain *et al*. 2016) sequences, and scaffolding using Hi-C data (Berkum *et al*. 2010). These new resources will enable genomic approaches to understanding the molecular basis of rapid adaptation in *C columbae*. More generally, the *C. columbae* genome provides comparative genomic data to understand the molecular basis of traits associated with parasitism that are shared among lice.

## Materials and Methods

### Animal tissue samples

All lice used in this study were drawn from natural populations infesting wild-caught feral rock pigeons (*Columba livia*) from Salt Lake City, UT. We maintained a captive pigeon colony to provide a constant source of *Columbicola columbae* lice used for sequencing.

The colony consisted of 15-20 pigeons, each harboring their own population of lice.

We reduced the nucleotide heterozygosity of our colony by creating a partially inbred population of lice. Initially, a single pair of lice (1 male, 1 female) was arbitrarily drawn from the pigeon colony and allowed to reproduce on a new, louse-free feral pigeon. After a period of 21 days, all immature lice were removed from the pigeon using CO_2_. At this point, these F1 lice are all full siblings. All offspring were then individually placed in glass vials stuffed with pigeon feathers to mature. Rearing lice individually in vials insured that F1s could not mate. Once mature, a single pair of unmated F1 adults (1 male, 1 female) were arbitrarily chosen and placed on a new, louse-free feral pigeon to mate and reproduce. Thus, all offspring on this new pigeon were the product of full-sibling mating and represented the first generation of inbreeding. These methods were repeated for eight generations.

After eight rounds of full-sibling inbreeding, all partially inbred lice were transferred to a new louse-free pigeon and left to mature and produce offspring. We left the lice on this pigeon for four months, which allowed the population to grow large enough to provide sufficient numbers for sequencing. The lice used for Illumina genomic DNA sequencing are derived from this partially inbred population. Reduced heterozygosity should result in higher quality polishing of the Oxford Nanopore-derived contigs with our Illumina data (see below). We pooled 100 adult lice for Illumina genomic DNA sequencing, 2000 adult lice for Oxford Nanopore genomic DNA sequencing, and 100 adult lice for Illumina RNA sequencing. All lice were drawn from the same partially inbred laboratory population, except for the lice used Oxford Nanopore sequencing, which were drawn from the main laboratory population from which the inbred population was derived. In addition, we generated Oxford Nanopore RNA sequencing reads from four different life stages of lice from the same starting population (100 lice each from larval instars 1, 2, and 3, and adults).

### Isolation of genetic material

DNA was isolated by grinding with a disposable homogenizer pestle (VWR, Radnor PA, USA) on ice for 30 minutes followed by DNA extraction with the Qiagen DNeasy extraction kit (Qiagen, Venlo, Netherlands). DNA for long read sequencing was extracted using the Qiagen DNA Blood and Tissue Midi kit. RNA was isolated using the Qiagen Oligotex mRNA mini kit.

### Illumina genomic DNA and RNA sequencing

Illumina DNA sequencing was performed using an Illumina HiSeq 2500 sequencer at the University of Utah High Throughput Genomics Core. We generated four libraries with mean insert sizes of 180 bp, 500 bp, 3500 bp, and 8200 bp. Genomic DNA was sequenced with paired-end 125-bp reads. cDNA sequencing was also performed on the Illumina HiSeq 2500 sequencer, producing paired end reads with a read length of 125bp.

### Oxford Nanopore genomic DNA and RNA sequencing

We generated long read genomic data using Oxford Nanopore MinION sequencers and a custom library preparation designed to increase read length. This protocol followed the standard procedure for producing 1d2 reads with kit LSK308 (Oxford Nanopore community, https://community.nanoporetech.com/protocols/), with the following modifications: (1) During all alcohol washes of magnetic SPRI beads, an additional wash was performed using Tris-EDTA to remove small DNA fragments. This step was performed quickly and without disturbing the beads to avoid dissolving all available DNA into solution. (2) All elutions from magnetic SPRI beads were performed after an incubation in elution buffer at 37°for 30 minutes. These practices improve the length of Oxford Nanopore sequencing reads (Urban *et al*. 2015).

We generated long mRNA reads using Oxford Nanopore MinION sequencers and a standard cDNA PCR-based sequencing method (PCS109, Oxford Nanopore community, https://community.nanoporetech.com/protocols/).

### Genome size estimation

We used the following formula (Liu *et al*. 2020) to estimate genome size from 21-mers counted from the Illumina sequencing data using *jellyfish* (Marçais and Kingsford 2011):

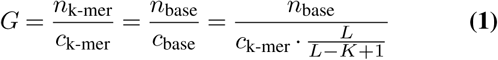

where *G* is the genome size, *n* is the total number of sequenced bases, *c* is the expected sequence coverage depth, *L* is the average sequencing read length, and *K* is the k-mer length.

### Genome assembly

We used *Trimmomatic* version 0.36 (Bolger *et al*. 2014) to trim Illumina input reads using the following settings: ILLUMINACLIP:adapters.fa:2:30:10 LEADING:20 TRAILING:20 MINLEN:30 CROP:85. then used *fastq-join* from *ea-utils* version 1.1.2-537 (Aronesty 2011) to join all short reads into pair joined reads, and used these throughout the assembly process. We used *Canu* v1.6 (Koren *et al*. 2017) with the parameter `genomeSize=220m` to assemble Oxford Nanopore genomic DNA reads, then polished the assembled contigs using *pilon* v1.22 (Walker *et al*. 2014) and the Illumina genomic DNA reads. The *pilon* software was run with the following switches: `-changes -vcf -vcfqe -tracks -fix all`.

The polished draft assembly was scaffolded by Phase Genomics using their proprietary scaffolding software (Burton *et al*. 2013; Peichel *et al*. 2017; Bickhart *et al*. 2017). We supplied Phase Genomics with approximately 1600 lice preserved at −80° for high molecular weight DNA extraction, Hi-C library preparation, and sequencing (Belton *et al*. 2012).

### Transcript selection and assembly

Standardized pipelines do not yet exist for selecting transcripts from raw Oxford Nanopore RNAseq reads. Therefore, we produced a custom pipeline that identifies putatively full-length transcripts to serve as evidence for genome annotation. In short, we aligned all RNAseq reads using *Minimap* (Li 2018), then clustered these alignments into sets that represent a gene using *Carnac-LR* (Marchet *et al*. 2019). We wrote a program, *wavy_choose*, that extracts the aligned reads from the original data, then identifies reads that likely represented full-length transcripts using *scipy*’s function scipy. signal. find_peaks_cwt () (Du *et al*. 2006) (Fig. 2). A more detailed version of the pipeline follows, below.

**Fig. 2.**
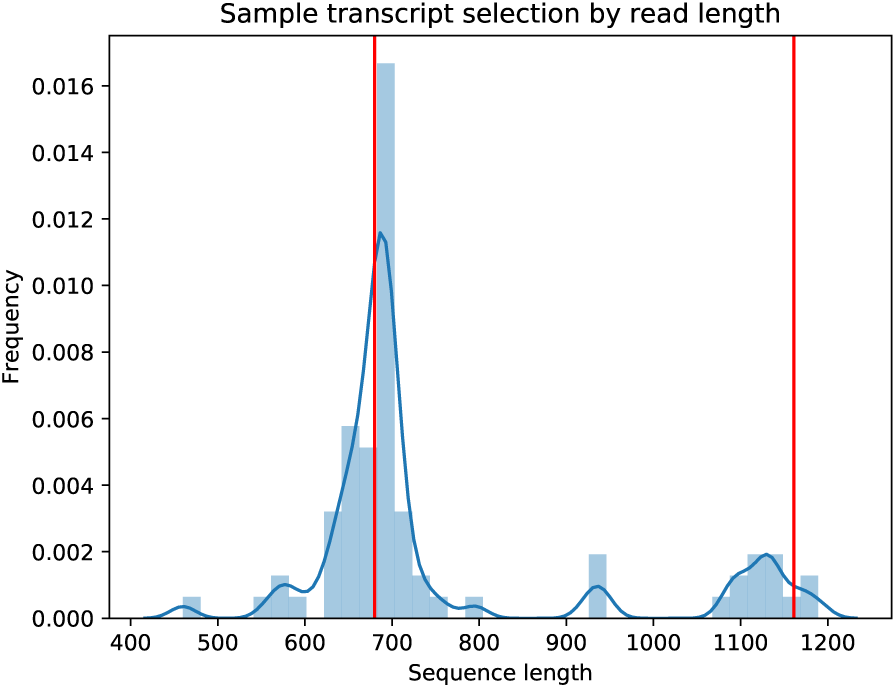
*wavy_choose* identifies likely full-length transcripts from clustered Oxford Nanopore reads. Depicted here is a histogram of read lengths (blue) for one *carnac-LR*-clustered set of reads. *wavy_choose* is able to identify two length peaks (red lines) in this transcript set, and discards all reads of other lengths. This process simplifies the transcriptome evidence dataset for MAKER, which uses the identified reads for gene annotation.

*Minimap* aligns long, low quality reads against one another, and can do so in an all-by-all comparison. *Carnac-LR* then clusters these reads into groups according to their alignments. Each *Carnac-LR*-clustered group of mRNA sequencing reads should represent all of the reads associated with a single gene, but if a gene has multiple alternative transcripts, *Carnac-LR* will not distinguish between them. The custom tool *wavy_choose* takes all of the clustered reads identified by *Carnac-LR* and identifies clusters within clusters that are most similar in both length and sequence. Because Oxford Nanopore reads are generally long enough to span an entire mRNA transcript, *wavy_choose* identifies the reads most likely to be complete transcripts by identifying the most common read lengths. It then removes all non-full-length reads from the analysis. This tool is especially well suited to transcript discovery, as multiple alternative transcripts may be identified from a single cluster of reads with overlapping sequence, and *wavy_choose* makes no assumptions as to the number of transcripts to identify.

The function find_peaks_cwt () uses continuous wavelet transformation, a technique from signal processing (Grossmann and Morlet 1984) to identify peaks in a 2-dimensional dataset. It does this by first convolving (transforming) the dataset to amplify the portion of the dataset that matches a wavelet with specified parameters (here, the default “Mexican hat” wavelet) and dampens the portions of the dataset that do not match the wavelet. The program then identifies local relative maxima that appear at the specified peak widths (here, 50 to 200 bp) and have sufficiently high signal-to-noise ratio (here 1.0). This widely accepted technique is straight-forward to apply in this context, but it is limited to detecting transcripts that have unique lengths. Two alternative transcripts of matching lengths would appear as a single peak in the length histogram. In these cases, reads from both alternative transcripts were retained in the final dataset. We kept at least one read per cluster of reads.

Untrimmed Illumina cDNA reads were assembled using *Trinity* using the*−−* jaccard_clip setting (Grabherr *et al*. 2011).

### Genome annotation

We used a combination of *wavy_choose*-selected Oxford Nanopore-derived transcripts, Illumina RNAseq-derived *Trinity* assemblies, and orthology information from Swissprot as evidence for gene models in *MAKER* (Cantarel *et al*. 2008), a widely used genome annotation tool. We used *AUGUSTUS* 3.3.1 to perform the gene finding portion of the *MAKER* pipeline. *BUSCO* (Simão *et al*. 2015) trains *AUGUSTUS* as part of the *BUSCO* pipeline, so we ran *BUSCO* on the genome assembly and used its *AUGUSTUS* training model during gene finding. We used both WU BLAST (Chao *et al*. 1992) and InterProScan (Jones *et al*. 2014) to match genes to their orthologs in the Uniprot-Swissprot database, and to provide the GO terms associated with genes in the final annotation set.

### Feature density analysis

We used *bedops* (Neph *et al*. 2012) to generate a .bed file of sliding windows across all chromosomes, then used *bedmap* (Neph *et al*. 2012) to count genes and repetitive sequences in these windows. Sliding windows were 1 Mb in width with a step length of 100 kb. For genes, we counted the total number of features identified by *MAKER* as “gene”s in its output .gff file. For repeats, we counted all *MAKER*-identified *repeatmasker* “match”es.

### Detection of bacterial contaminants

After assembly and annotation, we manually checked the louse genome for contamination with bacterial genomic sequences by identifying regions with unusually high gene density, *repeatmasker*-identified (Chen 2004) artifacts, and contiguous runs of bacterial genes. We also used *kraken* (Wood and Salzberg 2014) with the DustMasked MiniKraken DB database (https://ccb.jhu.edu/software/kraken/dl/minikraken_20171101_8GB_dustmasked.tgz) to identify known bacterial kmer contaminants.

We identified two sections of the genome that likely contained bacterial contamination, and removed them from the final assembly. The first section, at the beginning of chromosome 4, had a higher density of genes than any other region of the genome (280 genes per 10 kb, versus 64 genes per 10 kb in the bacteria-free genome). It also had a paucity of repetitive elements (262 repeats per 10 kb, as opposed to 800 elsewhere). *MAKER*’s annotation (see below) indicated that the majority of the region’s genes were bacterial in origin, and *BLASTn* searches (Zhang *et al*. 2000) against NCBI’s *nr* database (https://blast.ncbi.nlm.nih.gov/) confirmed this, as did *kraken*. The region also contained the annotation’s only instance of an explicit bacterial artifact identified by *repeatmasker*. The second region, on chromosome 8, was flagged as containing bacterial content by *kraken*. Both the chromosome 4 and 8 regions contained genes annotated by *MAKER* as similar to genes from the *Sodalis* clade, which contains the endosymbiont of the tsetse fly and a known bacterial endosymbiont of *C. columbae* (Fukatsu *et al*. 2007). 219 of the 554 genes in the chromosome 4 section are annotated as being *Sodalis*-related, as are 3 of the 4 genes in the chromosome 8 section. Thus, the totality of evidence led us to conclude that these regions on chromosomes 4 and 8 of our preliminary *C. columbae* genome assembly were bacterial contaminants from a known *Sodalis*-clade endosymbiont.

### Data availability

Raw sequence data for this project are publicly available through NCBI SRA (accession in progress). All analysis scripts are available through GitHub at https://github.com/jgbaldwinbrown/jgbutils. The genome assembly and annotation are available at NCBI GenBank (PRJNA662097).

## Results and Discussion

### Genome size estimation

We generated 2.92 ⨯ 10^10^ bases of genomic sequence using the Illumina short-read platform (mean read length after trimming = 107.2 bp). We estimated the genome size via *k*-mer counting (Liu *et al*. 2020) using *jellyfish* (Marçais and Kingsford 2011) (Fig. (Fig. 3). Using a k-mer size of 21, we estimate the genome size of *C. columbae* to be 230 Mb, within the range expected for insects.

**Fig. 3.**
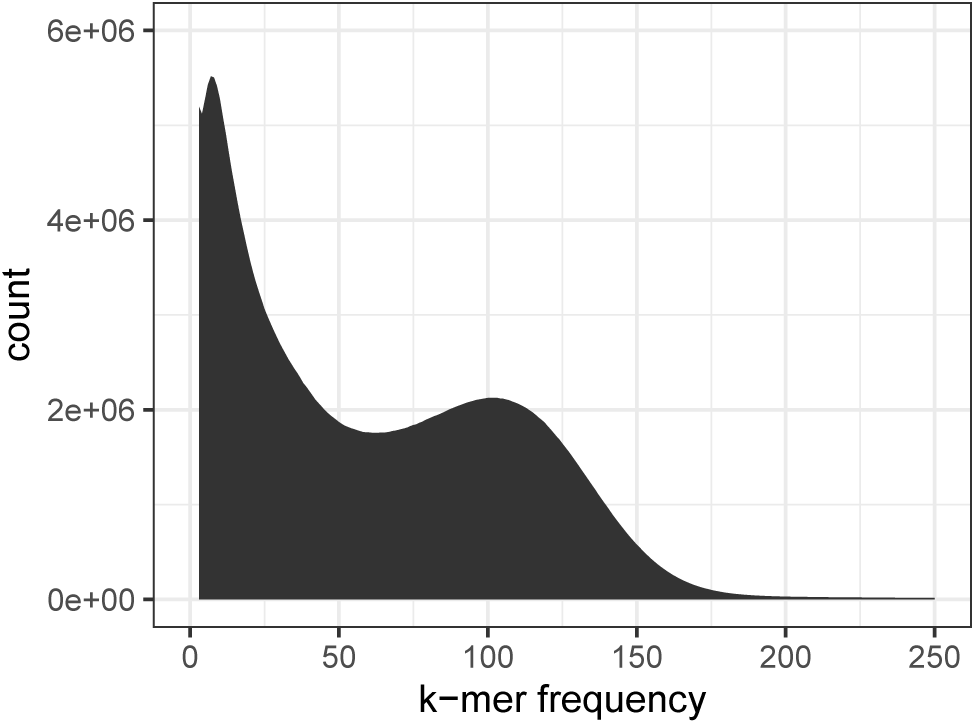
*jellyfish*-derived (Marçais and Kingsford 2011) 21-mer histogram based on Illumina reads from the *C. columbae* genome.

### Genome assembly and annotation summary

We generated a high-quality draft genome assembly using a combination of Illumina and Oxford Nanopore sequencing data, and Hi-C scaffolding. Our initial, unscaffolded assembly with *Canu* consists of 1193 contigs with a total length of 206 Mb, and an N50 contig length of 511 kb. We scaffolded the assembly using Hi-C data, producing chromosome-size scaffolds from the initial contigs. The final assembly comprises 12 chromosome-sized scaffolds and 380 small scaffolds, totaling 208 Mb of sequence. The N50 scaffold length for the final assembly is 17.7 Mb. Karyotyping evidence (Ries 1932) indicates that the *C. columbae* genome consists of 12 holocentric chromosomes. Based on this physical evidence, and the striking difference in size between the 12 largest scaffolds and all other scaffolds in the assembly (Fig. 4), we predict that each of the 12 scaffolds in the assembly represents one of the 12 karyotyped chromosomes.

**Fig. 4.**
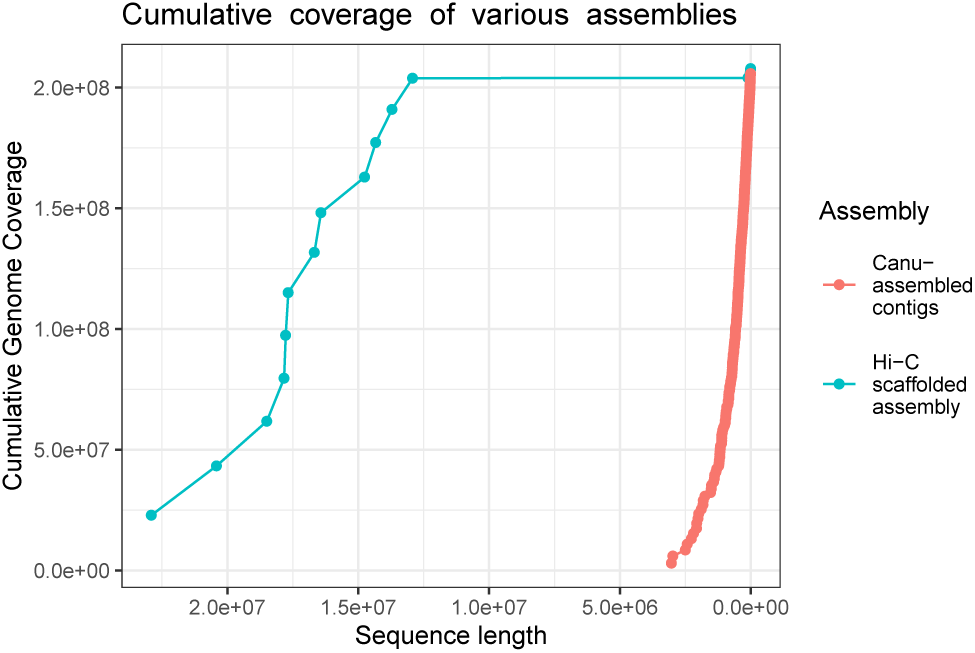
Cumulative coverage of initial and final (scaffolded) *C. columbae* genome assemblies, illustrating the improvement in contiguity by scaffolding with Hi-C data. All scaffolds in the assembly are plotted largest to smallest, from left to right. The x-axis indicates cumulative length of an assembly, and the y-axis corresponds to the cumulative portion of the genome covered by initial contigs (red dots) and final scaffolds (blue dots).

### Annotation

We annotated the genome using the *MAKER* pipeline, with transcriptome evidence from *Trinity*-assembled Illumina RNAseq reads and *wavy_choose*-selected Oxford Nanopore RNAseq reads (Fig. 2). We identified 19,139 transcripts from 13,362 genes. 13,246 of these genes are functionally annotated by BLAST using the Swissprot database, 8,354 are functionally annotated by similarity to InterPro or Pfam, and 13,248 are functionally annotated by either Swissprot, InterPro, or Pfam. *MAKER* produces a combined quality statistic called AED (Annotation edit distance; Eilbeck *et al*. (2009); Holt and Yandell (2011)). Perhaps owing to our use of long-read transcriptome sequencing, 10.3% of our annotated transcripts have ideal AED scores of 0 (Fig. 5), and only 5.6% of annotated transcripts have low-quality AED scores above 0.5. The abundance of low AED scores and relative dearth of low scores are indicators of a high-quality annotation.

**Fig. 5.**
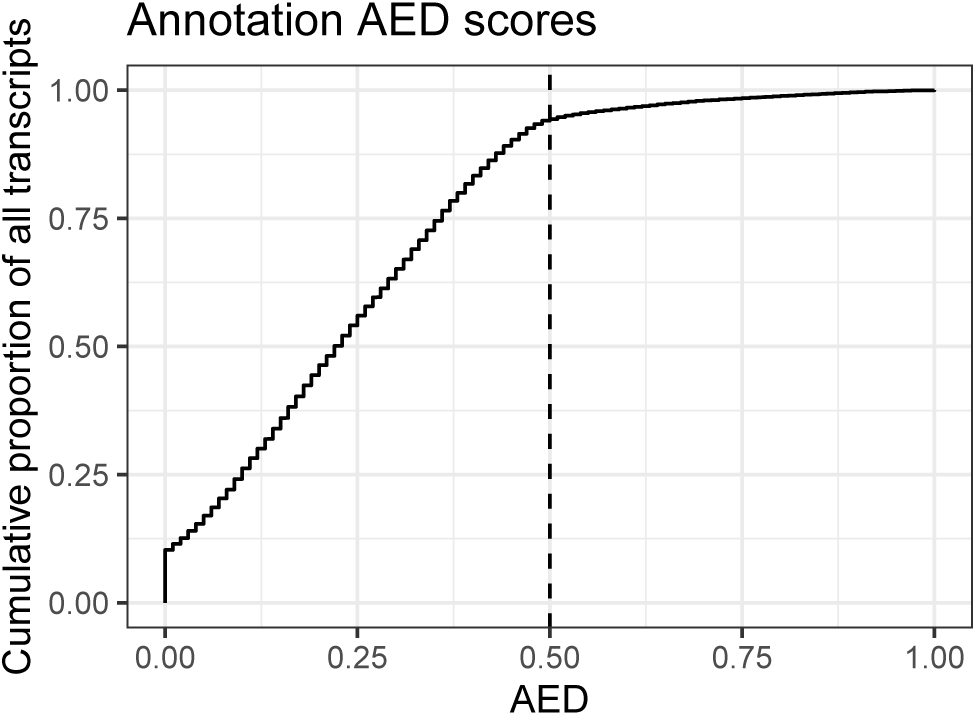
Cumulative annotation edit distance (AED) for all genes in the MAKER-derived annotation. 10.3% of genes have an AED of 0, while only 5.6% of genes have an AED above 0.5 (vertical dashed line).

### Genome completeness

We used BUSCO (Simão *et al*. 2015) to measure completeness of the genome by counting the number of highly conserved, single copy genes that should be present in insects. (Sup. Table 2). The reference genome, transcriptome, and translated transcriptome contain complete copies of 96%, 90%, and 87% of insect BUSCOs, respectively. These high values indicate that the *C. columbae* genome assembly is sufficiently complete for downstream comparative genomic analyses.

**Table 1.**
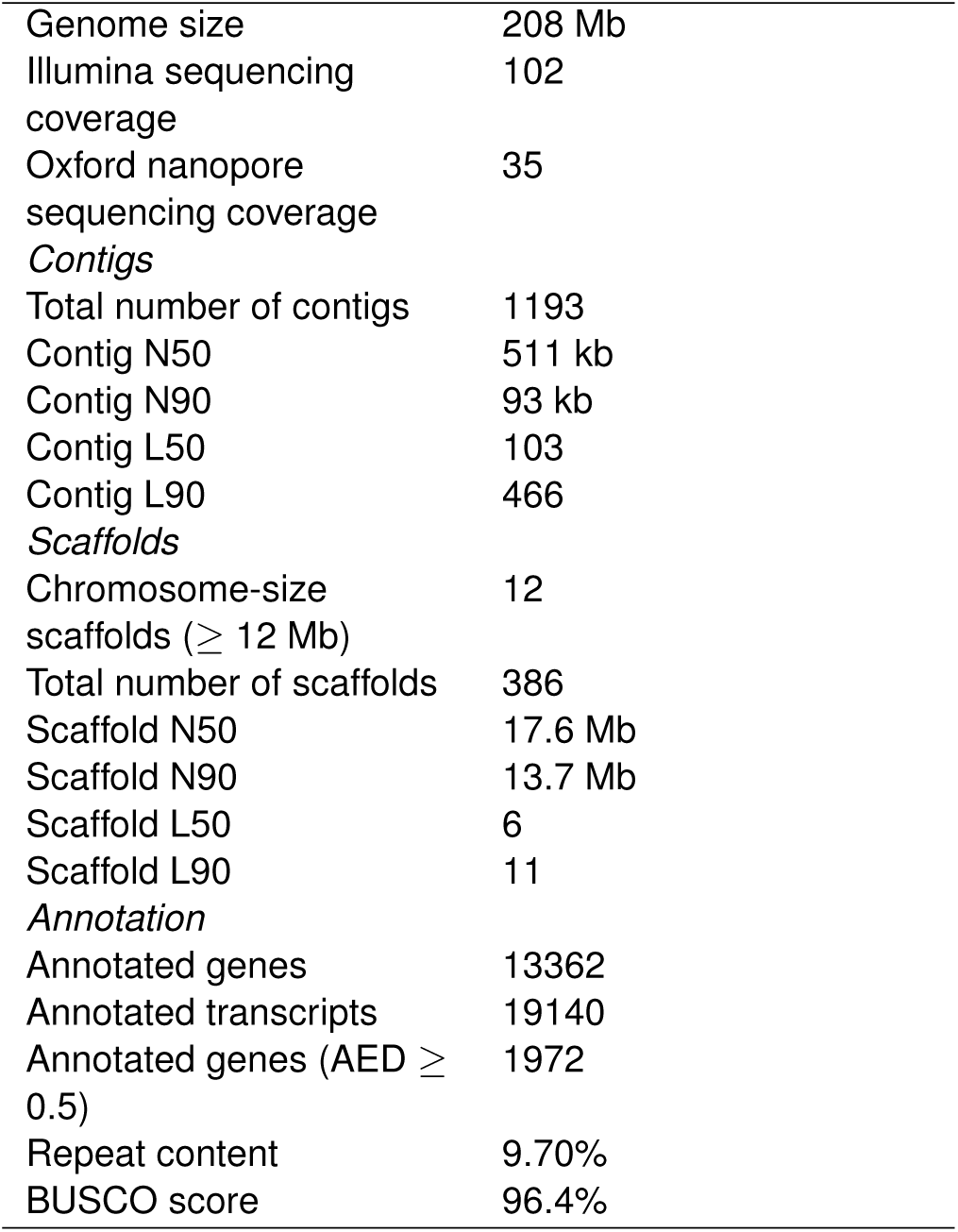
Assembly and annotation statistics.

|  |  |
| --- | --- |
| Genome size | 208 Mb |
| Illumina sequencing coverage | 102 |
| Oxford nanopore sequencing coverage | 35 |
| *Contigs* |  |
| Total number of contigs | 1193 |
| Contig N50 | 511 kb |
| Contig N90 | 93 kb |
| Contig L50 | 103 |
| Contig L90 | 466 |
| *Scaffolds* |  |
| Chromosome-size scaffolds ($\geq 12$ Mb) | 12 |
| Total number of scaffolds | 386 |
| Scaffold N50 | 17.6 Mb |
| Scaffold N90 | 13.7 Mb |
| Scaffold L50 | 6 |
| Scaffold L90 | 11 |
| *Annotation* |  |
| Annotated genes | 13362 |
| Annotated transcripts | 19140 |
| Annotated genes (AED $\geq 0.5$) | 1972 |
| Repeat content | 9.70% |
| BUSCO score | 96.4% |

**Table 2.** BUSCO results for genome completeness for the reference genome assembly, the annotated transcriptome, and the predicted proteome.

| Count | Genome | Transcriptome | Proteome |
| --- | --- | --- | --- |
| Total BUSCO groups searched | 1599 | 1490 | 1450 |
| Complete, single-copy BUSCOs | 1593 | 1440 | 1438 |
| Complete, duplicated BUSCOs | 6 | 50 | 12 |
| Fragmented BUSCOs | 25 | 54 | 55 |
| Missing BUSCOs | 34 | 114 | 153 |
| Complete BUSCOs (%) | 96.44 | 89.86 | 87.45 |
| Complete, single-copy BUSCOs (%) | 96.07 | 86.85 | 86.73 |
| Complete, duplicated BUSCOs (%) | 0.36 | 3.01 | 0.72 |
| Fragmented BUSCOs (%) | 1.50 | 3.25 | 3.31 |
| Missing BUSCOs (%) | 2.05 | 6.87 | 9.22 |
| Total BUSCO groups searched | 1658 | 1658 | 1658 |

### Repetitive elements

*Repeatmasker* identified 20.2 Mb (9.70%) of the genome as repetitive content. Of this 20.2 Mb, 65.8% is DNA transposons, 14.8% is LINEs, 8.6% is simple repeats, and 5.7% is LTR transposons (Wicker *et al*. 2007). The remainder (5.1%) is an assortment of transposable elements, low complexity regions, and satellites (Sup. table 3).

**Table 3.** Repetitive elements in the *C. columbae* genome.

| Identity | Number of bases | Percent of all bases | Percent of repetitive elements |
| --- | --- | --- | --- |
| DNA | 13283184 | 6.39 | 65.9 |
| LINE | 2980785 | 1.43 | 14.8 |
| Low_complexity | 506296 | 0.244 | 2.51 |
| LTR | 1156688 | 0.556 | 5.74 |
| Other | 684 | 0.000329 | 0.00339 |
| RC | 126151 | 0.0607 | 0.626 |
| Retroposon | 749 | 0.000360 | 0.00371 |
| RNA | 16720 | 0.00804 | 0.0829 |
| rRNA | 33934 | 0.0163 | 0.168 |
| Satellite | 48279 | 0.0232 | 0.239 |
| Simple_repeat | 1745924 | 0.840 | 8.660 |
| SINE | 112911 | 0.0543 | 0.560 |
| snRNA | 24864 | 0.0120 | 0.123 |
| tRNA | 54873 | 0.0264 | 0.272 |
| Unknown | 66922 | 0.0322 | 0.332 |

Repetitive content in the *C. columbae* genome is, therefore, considerably higher than in *P. humanus*. In the latter species, only 1% of the genome is annotated as class I (retrotransposons, including LTR, LINE, and SINE) or class II (DNA transposons) transposable elements, and 6.9% is tandem repeats (Kirkness *et al*. 2010).

Kirkness *et al*. (2010) predicted that the monophagous, permanently parasitic lifestyle of lice should lead to reduced genomes due to the reduced need to seek food and avoid enemies compared to free-living species. While Kirkness et al. identify a reduction in gene families related to sensing, their conclusion that overall genome size is affected by lifestyle is not supported by the genome size of *C. columbae*, which has a genome size and number of genes that are more typical for a free-living insect. Indeed, both *C. columbae* and *P. humanus* appear to have a full complement of genes, and while *P. humanus* has a small genome and a reduction in transposable elements, *C. columbae* has neither of these. The pattern of reduced genome size and reduction in TE content without loss of genes is characteristic of high-population-size species (Lefébure *et al*. 2017). However, a robust estimate of the population size of *P. humanus*, combined with evidence ruling out alternative hypotheses, would be necessary to demonstrate that population size drove the reduced genome size in *P. humanus*. Other authors (Oliver *et al*. 2007) have hypothesized that large populations may not actually be under selection to have smaller genomes.

### Genomic evidence for the lack of centromeres

Centromeres are characterized by a depletion of genic content and an increase in repetitive content (Jain *et al*. 2018). Based on these criteria (Fig. 6), we find no evidence for centromeres in any of the *C. columbae* chromosomes. Presence of genes is moderately anti-correlated with presence of simple repetitive sequences (r = −0.28, 1 Mb sliding windows). Still, the overall repeat density is not correlated with gene density, and both measures are relatively consistent across the genome. Many chromosomes (c.f., Fig. 6, chromosomes 6 and 7) have a twin-peaked pattern of simple repeats, in which chromosome ends and centers have high genic content and low repeat content, but the genomic segments between the ends and the center have high repeat content and low genic content. It is possible that these twin peaks of simple repeat content are the centromeres in a polycentromeric chromosome, and that the chromosomes were actually misclassified as holocentric based on karyotyping evidence.

**Fig. 6.**
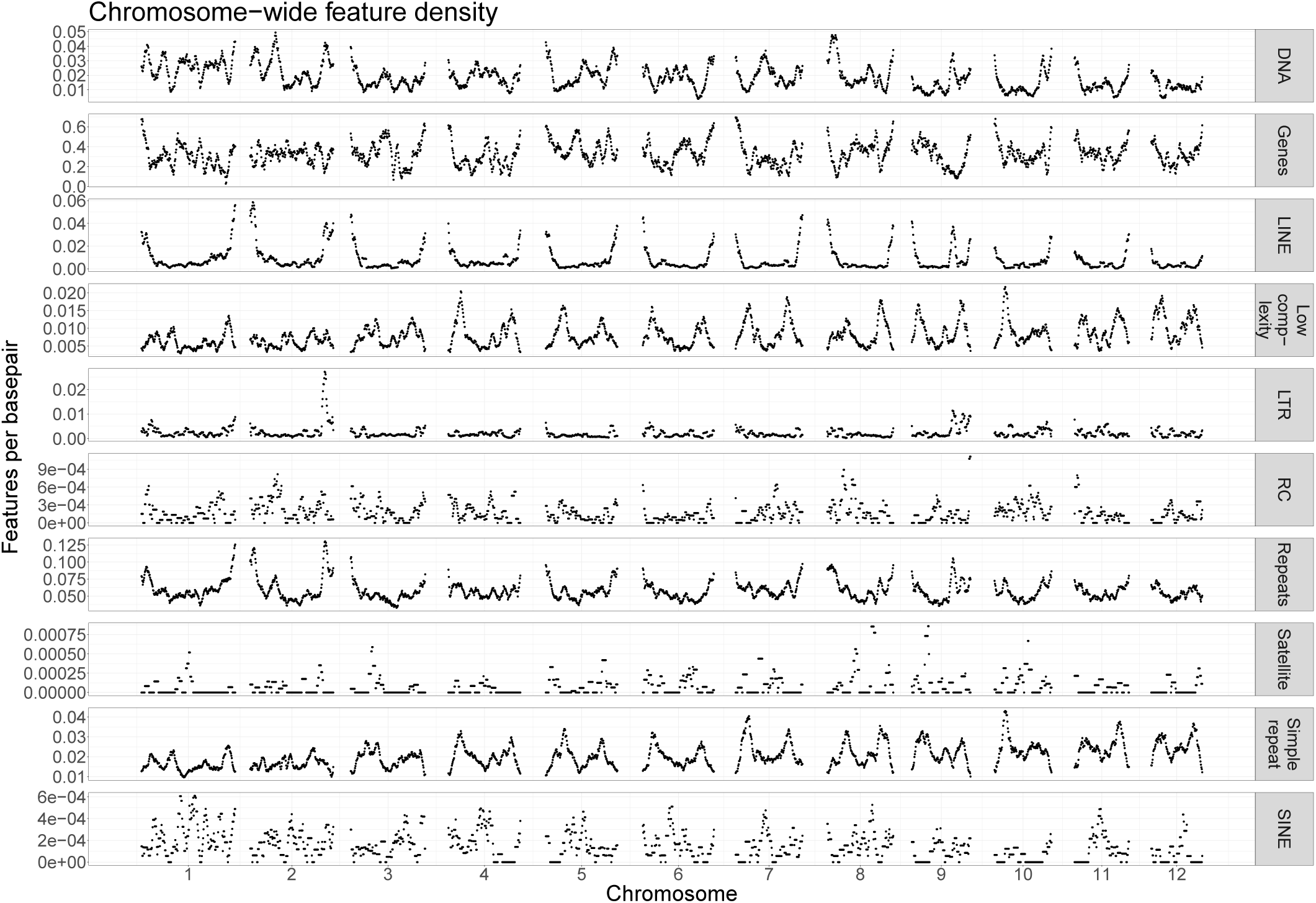
Gene and repeat density in 1Mb-wide sliding windows across the *C. columbae* genome show that there are no clear centromeres, and gene and simple repeat density are negatively correlated.

### Comparisons to the closest sequenced relative

The closest relative of *C. columbae* with an assembled genome is the human body louse *Pediculus humanus. C. columbae* and *P. humanus* are thought to have diverged 65 million years ago (Johnson *et al*. 2018). *P. humanus* has five metacentric chromosomes and one telocentric chromosome (Kirkness *et al*. 2010), in contrast to the twelve putatively holocentric chromosomes described here, and has a genome assembly size of 108 Mb, approximately half that of the 208-Mb *C. columbae* genome assembly. The *C. columbae* genome has a typical genome-wide GC content of 36%, while *P. humanus* has an extremely AT-rich genome with 28% GC content, making *C. columbae* the more typical insect genome of the two.

### Synteny analysis

We used the default settings of *SynIma* (Farrer 2017) to identify synteny between *C. columbae* and *P. humanus* (Fig. 7). We were unable to test for chromosome-scale syntenic blocks between *P. humanus* and *C. columbae* due to the low contiguity of the *P. humanus* genome. However, we found very few locations in which synteny is broken between a *P. humanus* scaffold and a *C. columbae* scaffold, showing that short-range synteny is almost entirely conserved between these species.

**Fig. 7.**
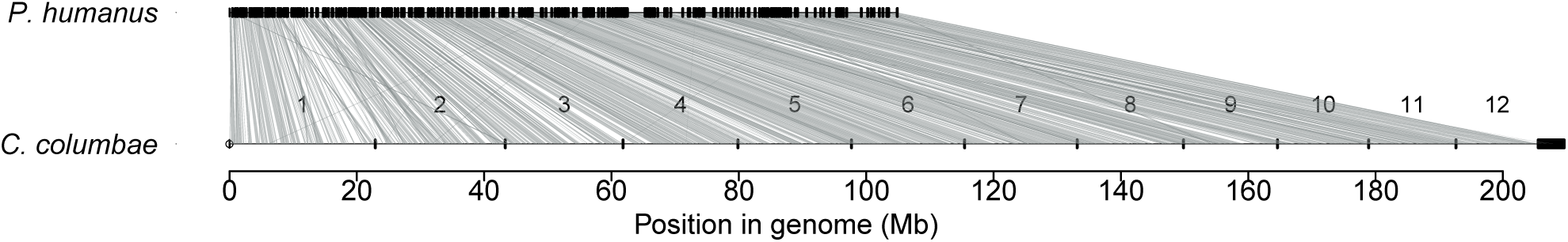
Short range synteny is largely conserved between *C. columbae* (bottom) and *P. humanus* (top) genomic scaffolds. Lines connecting scaffolds from each genome assembly represent the positions of orthologous genes. *P. humanus* contigs were aligned to the *C. columbae* genome in order and orientation using *SynIma*.

### Functional annotation reveals depletion of environmental sensing and metabolic genes

*P. humanus* has a small complement of opsins (3, as opposed to 275 in *D. melanogaster*) and G protein-coupled receptors (GPCR, 104, as opposed to 408 in *D. melanogaster*) (Kirkness *et al*. 2010; Thurmond *et al*. 2019). Similarly, we find that only 2 annotated genes in *C. columbae* are associated with opsin gene ontology term (GO:00007602) and only 107 genes are associated with GPCR GO category (GO:00004930). This reduced repertoire of sensory system genes supports the hypothesis that the relatively static environments encountered by lice and other ectoparasites relaxes selection on the ability to sense and respond to stimuli in more variable environments (Kirkness *et al*. 2010). *C. columbae* is incapable of surviving off of its obligate host, so there might be little selection to retain complex visual, olfactory, or other complex sensory acuity.

We find support for the hypothesis that specific gene families, such as those relating to sensory capabilities and metabolism, are reduced in obligate parasites (Jackson 2015).

*P. humanus* is massively depleted in terms of odorant receptors, gustatory receptors, and chemosensory proteins, and *C. columbae* shows the same pattern. For example, *C. columbae* has only 13 genes with olfactory receptor activity (GO:0004984) and *P. humanus* has only 10, compared with 152 in *D. melanogaster* (Thurmond *et al*. 2019). *C. columbae* has 2 genes associated with taste receptor activity (GO:0008527) and *P. humanus* has 6, yet *D. melanogaster* has 150. We speculate that this dramatic depletion of taste receptor genes is due to the homogeneous diet of ectoparasitic lice. The diet of *C. columbae*, for instance, consists entirely of pigeon feathers (Ash 1960).

Another highly depleted gene functional category in *P. humanus* is the insulin signaling / TOR pathway. Kirkness *et al*. (2010) show that the canonical pathway appears non-functional in *P. humanus*, with only one gene having *P. humanus* EST-derived evidence for its expression. BLAST evidence indicates that other TOR pathway genes are reduced to a single copy in *P. humanus*, including genes such as *dilps* and *eIF-4E* (class I), which respectively have 6 and 7 copies in *D. melanogaster* (Kirkness *et al*. 2010). We find the same qualitative result in *C. columbae*, with no annotated genes associated with the insulin receptor signaling pathway (GO:0008286). Finally, the complement of detoxification genes is depleted in both *P. humanus* and *C. columbae*, with *C. columbae* having no annotated genes associated with detoxification (GO:0098754).

The striking reduction in sensory and metabolic gene categories in *C. columbae* and *P. humanus* could be due to independent gene loss in each lineage, inheritance of a depleted repertoire from a common ancestor, or a combination of the two. Loss of the same suite of genes in each species would be consistent with inheritance of a reduced sensory repertoire from a common ancestor, while loss of different genes in each species would indicate independent reductions. Reciprocal best BLAST hits of *C. columbae* and *P. humanus* genes to a shared outgroup, *Drosophila melanogaster*, indicate that the identities of the lost and retained genes are mostly the same between the two louse species (Table 4), thereby supporting the hypothesis of ancestral loss. We note the possibility that these “missing” genes are not actually absent from the genomes of *C. columbae* and *P. humanus*, but are simply not annotated in their respective genomes. However, the BUSCO completeness score of 96.4% for the *C. columbae* genome renders large-scale incompleteness and misannotation less likely.

**Table 4.** Reciprocal best-hit BLAST of the proteomes of *C. columbae* and *P. humanus* against *D. melanogaster* reveals the identity of the retained genes in depleted gene families is largely the same in both species. The first column is the tested family of genes. The second column is the number of genes assigned the corresponding GO term in the *D. melanogaster* proteome. The third and fourth columns, respectively, are the numbers of reciprocal best BLAST hits with *D. melanogaster* genes by genes from either *C. columbae* or *P. humanus*. The fifth column is the number of reciprocal best BLAST hits that had the same *D. melanogaster*-derived identity when BLASTing against *C. columbae* or *P. humanus*.

| Gene family | *D. melanogaster* genes | *C. columbae* hits | *P. humanus* hits | Shared hits |
| --- | --- | --- | --- | --- |
| Opsin | 275 | 40 | 40 | 28 |
| GPCR | 408 | 66 | 69 | 50 |
| Olfactory receptor activity | 152 | 6 | 8 | 6 |
| Taste | 150 | 4 | 3 | 3 |
| Odorant binding | 248 | 6 | 7 | 4 |
| Insulin | 349 | 59 | 61 | 46 |
| Tor | 225 | 51 | 57 | 47 |
| Chemosensory behavior | 441 | 54 | 57 | 44 |
| Detoxification | 132 | 16 | 18 | 16 |

### Summary

In summary, we report a high-quality draft genome assembly and annotation for the ectoparasitic louse *Columbicola columbae*. We present a new method for selecting transcripts from long-read Oxford Nanopore RNA-seq data, and use the selected transcripts as evidence for the *MAKER* annotation pipeline. We find massive depletion of sensory and metabolic genes, similar to findings for the only other published louse genome. Comparative analysis points to loss of many of the same genes in both lice, suggesting that these genes were probably lost in a common ancestor at least 65 million years ago. Looking ahead, the *C. columbae* genome provides new tools for comparative genomics of lice and other insects, and poises us to understand the molecular basis of rapid evolution in *C. columbae* itself.

## Acknowledgements

This work was funded by the National Science Foundation dimensions in biodiversity grants NSF DEB-1342604 and DEB-1342600. We thank Juan C. Altuna for assistance in creating the louse inbreeding protocol and for maintenance of the captive populations of pigeons and lice. We gratefully acknowledge support and resources from the Center for High Performance Computing at the University of Utah. We also thank Mark Yandell for providing computational resources, and Carson Holt for technical advice and assistance with MAKER.

## Literature Cited

Adly, E., M. Nasser, D. Soliman, D. R. Gustafsson, and M. Shehata, 2019 New records of chewing lice (Phthiraptera: Amblycera, Ischnocera) from Egyptian pigeons and doves (Columbiformes), with description of one new species. Acta Tropica 190: 22–27.

Aronesty, E., 2011 ea-utils: Command-line tools for processing biological sequencing data. Durham, NC.

Ash, J. S., 1960 A study of the mallophaga of birds with particular reference to their ecology. Ibis 102: 93–110, _eprint: https://onlinelibrary.wiley.com/doi/pdf/10.1111/j.1474-919X.1960.tb05095.x.

Belton, J.-M., R. P. McCord, J. Gibcus, N. Naumova, Y. Zhan, et al., 2012 Hi-C: A comprehensive technique to capture the conformation of genomes. Methods (San Diego, Calif.) 58.

Bennett, S., 2004 Solexa Ltd. Pharmacogenomics 5: 433–438.

Berkum, N. L. v., E. Lieberman-Aiden, L. Williams, M. Imakaev, A. Gnirke, et al., 2010 Hi-C: A Method to Study the Three-dimensional Architecture of Genomes. JoVE (Journal of Visualized Experiments) p. e1869.

Bickhart, D. M., B. D. Rosen, S. Koren, B. L. Sayre, A. R. Hastie, et al., 2017 Single-molecule sequencing and chromatin conformation capture enable de novo reference assembly of the domestic goat genome. Nature Genetics 49: 643–650.

Bolger, A. M., M. Lohse, and B. Usadel, 2014 Trimmomatic: a flexible trimmer for Illumina sequence data. Bioinformatics 30: 2114–2120.

Burton, J. N., A. Adey, R. P. Patwardhan, R. Qiu, J. O. Kitzman, et al., 2013 Chromosome-scale scaffolding of de novo genome assemblies based on chromatin interactions. Nature Biotechnology 31: 1119–1125.

Bush, S., D. Kim, M. Reed, and D. Clayton, 2010 Evolution of cryptic coloration in ectoparasites. The American Naturalist 176: 529–535, Publisher: The University of Chicago Press.

Bush, S. E. and D. H. Clayton, 2006 The role of body size in host specificity: reciprocal transfer experiments with feather lice. Evolution 60: 2158–2167, _eprint: https://onlinelibrary.wiley.com/doi/pdf/10.1111/j.0014-3820.2006.tb01853.x.

Bush, S. E., R. D. Price, and D. H. Clayton, 2009 Descriptions of eight new species of feather lice in the genus Columbicola (Phthiraptera: Philopteridae), with a comprehensive world checklist. Journal of Parasitology 95: 286–294, Publisher: Allen Press.

Bush, S. E., E. Sohn, and D. H. Clayton, 2006 Ecomorphology of parasite attachment: experiments with feather lice. Journal of Parasitology 92: 25–31, Publisher: Allen Press.

Bush, S. E., S. M. Villa, J. C. Altuna, K. P. Johnson, M. D. Shapiro, et al., 2019 Host defense triggers rapid adaptive radiation in experimentally evolving parasites. Evolution Letters 3: 120–128.

Cantarel, B. L., I. Korf, S. M. C. Robb, G. Parra, E. Ross, et al., 2008 MAKER: An easy-to-use annotation pipeline designed for emerging model organism genomes. Genome Research 18: 188–196.

Chao, K.-M., W. R. Pearson, and W. Miller, 1992 Aligning two sequences within a specified diagonal band. Bioinformatics 8: 481–487, Publisher: Oxford Academic.

Chen, N., 2004 Using RepeatMasker to identify repetitive elements in genomic sequences. Current Protocols in Bioinformatics 5: 4.10.1–4.10.14.

Clayton, D. H., 1990 Mate choice in experimentally parasitized rock doves: lousy males lose. Integrative and Comparative Biology 30: 251–262, Publisher: Oxford Academic.

Clayton, D. H., 1991 Coevolution of avian grooming and ectoparasite avoidance. Bird–Parasite Interactions: Ecology, Evolution and Behaviour 14: 258–289, Publisher: Oxford University Press Oxford, UK.

Clayton, D. H., R. J. Adams, and S. E. Bush, 2008 Phthiraptera, the chewing lice. Parasitic Diseases of Wild Birds pp. 515–526, Publisher: Wiley Online Library.

Clayton, D. H., S. E. Bush, B. M. Goates, and K. P. Johnson, 2003 Host defense reinforces host–parasite cospeciation. Proceedings of the National Academy of Sciences 100: 15694–15699, Publisher: National Academy of Sciences Section: Biological Sciences.

Clayton, D. H., S. E. Bush, and K. P. Johnson, 2015 Coevolution of life on hosts: integrating ecology and history. University of Chicago Press, Google-Books-ID: z4gCCwAAQBAJ.

Clayton, D. H., P. L. M. Lee, D. M. Tompkins, and E. D. Brodie III, 1999 Reciprocal natural selection on hostparasite phenotypes. The American Naturalist 154: 261–270, Publisher: The University of Chicago Press.

Clayton, D. H. and D. M. Tompkins, 1995 Comparative effects of mites and lice on the reproductive success of rock doves (Columba livia). Parasitology 110: 195–206.

Darwin, C., 1868 The Variation of Animals and Plants Under Domestication, volume 1. Orange Judd & Co, New York, NY.

de Meeûs, T. and F. Renaud, 2002 Parasites within the new phylogeny of eukaryotes. Trends in Parasitology 18: 247–251.

Du, P., W. A. Kibbe, and S. M. Lin, 2006 Improved peak detection in mass spectrum by incorporating continuous wavelet transform-based pattern matching. Bioinformatics 22: 2059–2065, Publisher: Oxford Academic.

Durden, L. A. and G. G. Musser, 1994 The sucking lice (Insecta, Anoplura) of the world: a taxonomic checklist with records of mammalian hosts and geographical distributions. Bullentin of the American Museum of Natural History 218, Accepted: 2005-11-22T22:50:41Z Publisher: [New York] : American Museum of Natural History.

Eilbeck, K., B. Moore, C. Holt, and M. Yandell, 2009 Quantitative measures for the management and comparison of annotated genomes. BMC Bioinformatics 10: 67.

Farrer, R. A., 2017 Synima: a Synteny imaging tool for annotated genome assemblies. BMC Bioinformatics 18: 507.

Fukatsu, T., R. Koga, W. A. Smith, K. Tanaka, N. Nikoh, et al., 2007 Bacterial endosymbiont of the slender pigeon louse, Columbicola columbae, allied to endosymbionts of grain weevils and tsetse flies. Applied and Environmental Microbiology 73: 6660–6668.

Gibbs, D., E. Barnes, and J. Cox, 2001 Pigeons and Doves: a Guide to the Pigeons and Doves of the World, volume 13. A&C Black, London.

Grabherr, M. G., B. J. Haas, M. Yassour, J. Z. Levin, D. A. Thompson, et al., 2011 Trinity: reconstructing a fulllength transcriptome without a genome from RNA-Seq data. Nature Biotechnology 29: 644–652.

Grossmann, A. and J. Morlet, 1984 Decomposition of hardy functions into square Integrable wavelets of constant shape. SIAM Journal on Mathematical Analysis 15: 723–736, Publisher: Society for Industrial and Applied Mathematics.

Gustafsson, D. R., M. Tsurumi, and S. E. Bush, 2015 The chewing lice (Insecta: Phthiraptera: Ischnocera: Amblycera) of Japanese pigeons and doves (Columbiformes), with descriptions of three new species. Journal of Parasitology 101: 304–313, Publisher: Allen Press.

Harbison, C. W. and D. H. Clayton, 2011 Community interactions govern host-switching with implications for host–parasite coevolutionary history. Proceedings of the National Academy of Sciences 108: 9525–9529, Publisher: National Academy of Sciences Section: Biological Sciences.

Holt, C. and M. Yandell, 2011 MAKER2: an annotation pipeline and genome-database management tool for second-generation genome projects. BMC Bioinformatics 12: 491.

Jackson, A. P., 2015 The evolution of parasite genomes and the origins of parasitism. Parasitology 142: S1–S5.

Jain, M., H. E. Olsen, B. Paten, and M. Akeson, 2016 The Oxford Nanopore MinION: delivery of nanopore sequencing to the genomics community. Genome Biology 17: 239.

Jain, M., H. E. Olsen, D. J. Turner, D. Stoddart, K. V. Bulazel, et al., 2018 Linear assembly of a human centromere on the Y chromosome. Nature Biotechnology 36: 321–323, Number: 4 Publisher: Nature Publishing Group.

Johnson, K. P., J. R. Malenke, and D. H. Clayton, 2009 Competition promotes the evolution of host generalists in obligate parasites. Proceedings of the Royal Society B: Biological Sciences 276: 3921–3926.

Johnson, K. P., N.-p. Nguyen, A. D. Sweet, B. M. Boyd, T. Warnow, et al., 2018 Simultaneous radiation of bird and mammal lice following the K-Pg boundary. Biology Letters 14: 20180141, Publisher: Royal Society.

Johnson, K. P., D. L. Reed, S. L. Hammond Parker, D. Kim, and D. H. Clayton, 2007 Phylogenetic analysis of nuclear and mitochondrial genes supports species groups for Columbicola (Insecta: Phthiraptera). Molecular Phylogenetics and Evolution 45: 506–518.

Jones, P., D. Binns, H.-Y. Chang, M. Fraser, W. Li, et al., 2014 InterProScan 5: genome-scale protein function classification. Bioinformatics 30: 1236–1240.

Kirkness, E. F., B. J. Haas, W. Sun, H. R. Braig, M. A. Perotti, et al., 2010 Genome sequences of the human body louse and its primary endosymbiont provide insights into the permanent parasitic lifestyle. Proceedings of the National Academy of Sciences 107: 12168–12173, Publisher: National Academy of Sciences Section: Biological Sciences.

Koren, S., B. P. Walenz, K. Berlin, J. R. Miller, N. H. Bergman, et al., 2017 Canu: scalable and accurate longread assembly via adaptive k-mer weighting and repeat separation. Genome Research 27: 722–736.

Lefébure, T., C. Morvan, F. Malard, C. François, L. Konecny-Dupré, et al., 2017 Less effective selection leads to larger genomes. Genome Research 27: 1016–1028.

Li, H., 2018 Minimap2: pairwise alignment for nucleotide sequences. Bioinformatics (Oxford, England) 34: 3094–3100.

Liu, B., Y. Shi, J. Yuan, X. Hu, H. Zhang, et al., 2020 Estimation of genomic characteristics by analyzing k-mer frequency in de novo genome projects. 1308.2012 [q-bio] 1308.2012.

Marchet, C., L. Lecompte, C. D. Silva, C. Cruaud, J.-M. Aury, et al., 2019 De novo clustering of long reads by gene from transcriptomics data. Nucleic Acids Research 47: e2.#x2013;e2, Publisher: Oxford Academic.

Marshall, A. G., 1981 The Ecology of Ectoparasitic Insects. Academic Press, Cambridge, Massachusetts.

Martin, M., 1934 Life history and habits of the pigeon louse (Columbicola columbae [Linnaeus]). The Canadian Entomologist 66: 6–16, Publisher: Cambridge University Press.

Marçais, G. and C. Kingsford, 2011 A fast, lock-free approach for efficient parallel counting of occurrences of k-mers. Bioinformatics 27: 764–770.

Mullen, G. R. and L. A. Durden, 2009 Medical and Veterinary Entomology. Academic Press, Cambridge, Massachusetts, Google-Books-ID: T8CWvVGwKhoC.

Nelson, B. C. and M. D. Murray, 1971 The distribution of mallophaga on the domestic pigeon (Columba livia). International Journal for Parasitology 1: 21–29.

Neph, S., M. S. Kuehn, A. P. Reynolds, E. Haugen, R. E. Thurman, et al., 2012 BEDOPS: high-performance genomic feature operations. Bioinformatics 28: 1919–1920.

Oliver, M. J., D. Petrov, D. Ackerly, P. Falkowski, and O. M. Schofield, 2007 The mode and tempo of genome size evolution in eukaryotes. Genome Research 17: 594–601.

Peichel, C. L., S. T. Sullivan, I. Liachko, and M. A. White, 2017 Improvement of the threespine stickleback genome using a Hi-C-based proximity-guided assembly. Journal of Heredity 108: 693–700.

Rakshpal, R., 1959 On the behaviour of pigeon louse, Columbicola columbae Linn. (Mallophaga). Parasitology 49: 232–241, Publisher: Cambridge University Press.

Ries, E., 1932 Die prozesse der eibildung und des eiwachstums bei pediculiden und mallophagen. Zeitschrift für Zellforschung und Mikroskopische Anatomie 16: 314–388.

Rudolph, D., 1983 The water-vapour uptake system of the phthiraptera. Journal of Insect Physiology 29: 15–25.

Shapiro, M. D. and E. T. Domyan, 2013 Domestic pigeons. Current Biology 23: R302–R303, Publisher: Elsevier.

Simão, F. A., R. M. Waterhouse, P. Ioannidis, E. V. Kriventseva, and E. M. Zdobnov, 2015 BUSCO: assessing genome assembly and annotation completeness with single-copy orthologs. Bioinformatics 31: 3210–3212.

Smith, V. S., 2004 The chewing lice: world checklist and biological overview. Systematic Biology 53: 666–668, Publisher: Oxford Academic.

Stenram, H., 1956 The ecology of Columbicola columbae L.(Mallophaga). Opusculata Entomologica 21: 170–190.

Thurmond, J., J. L. Goodman, V. B. Strelets, H. Attrill, L. S. Gramates, et al., 2019 FlyBase 2.0: the next generation. Nucleic Acids Research 47: D759–D765, Publisher: Oxford Academic.

Urban, J. M., J. Bliss, C. E. Lawrence, and S. A. Gerbi, 2015 Sequencing ultra-long DNA molecules with the Oxford Nanopore MinION. bioRxiv p. 019281, Publisher: Cold Spring Harbor Laboratory Section: New Results.

Villa, S. M., J. C. Altuna, J. S. Ruff, A. B. Beach, L. I. Mulvey, et al., 2019 Rapid experimental evolution of reproductive isolation from a single natural population. Proceedings of the National Academy of Sciences 116: 13440–13445.

Walker, B. J., T. Abeel, T. Shea, M. Priest, A. Abouelliel, et al., 2014 Pilon: an integrated tool for comprehensive microbial variant detection and genome assembly improvement. PLOS ONE 9: e112963.

Wicker, T., F. Sabot, A. Hua-Van, J. L. Bennetzen, P. Capy, et al., 2007 A unified classification system for eukaryotic transposable elements. Nature Reviews Genetics 8: 973–982, Number: 12 Publisher: Nature Publishing Group.

Wood, D. E. and S. L. Salzberg, 2014 Kraken: ultrafast metagenomic sequence classification using exact alignments. Genome Biology 15: R46.

Zhang, Z., S. Schwartz, L. Wagner, and W. Miller, 2000 A greedy algorithm for aligning DNA sequences. Journal of Computational Biology 7: 203–214, Publisher: Mary Ann Liebert, Inc., publishers.

